# High-Throughput Characterization of Trends in Transmembrane Helix Partitioning into Membrane Domains

**DOI:** 10.64898/2026.05.14.725159

**Authors:** Jonathan Thelen, Melanie König, Maisa Vuorte, Jarkko Liimatainen, Matti Javanainen, Fabio Lolicato

## Abstract

The plasma membrane is a laterally heterogeneous environment in which lipid organization plays a central role in regulating protein function. In model systems, this heterogeneity is often described in terms of coexisting liquid-ordered (L_o_) and liquid-disordered (L_d_) phases, commonly associated with the lipid raft concept. Despite extensive experimental and computational efforts, the molecular determinants governing protein partitioning between these domains remain poorly understood, largely due to the limited number of systems studied.

Here, we address this challenge using a high-throughput computational approach, systematically analyzing the partitioning behavior of almost 5,000 helical transmembrane peptides in phase-separating lipid membranes. Across all simulations, we find that none of the peptides exhibit a clear preference for the L_o_ phase, while the vast majority partition into the L_d_ phase. This observation is consistent with experimental results in simplified membrane systems and suggests that commonly used ternary lipid mixtures may not fully capture the physicochemical environment governing protein sorting in biological membranes. In addition, we identify a subset of peptides that preferentially localize at the L_o_/L_d_ interface. These interfacial peptides display distinct sequence characteristics, indicating that boundary localization is governed by specific combinations of residue composition and spatial arrangement rather than a single dominant feature. Overall, our results reveal that transmembrane helix partitioning in model membranes is dominated by a preference for disordered environments, with interfacial localization emerging as a distinct and potentially functional behavior.

## Introduction

The plasma membrane exhibits pronounced lateral heterogeneity, organizing membrane proteins into distinct environments that regulate and support their functions. The lipid raft concept was introduced to describe this organization, proposing small, dynamic, cholesterol- and sphingolipid-rich domains whose size, lifetime, and functional relevance remain under debate.^1–7^ In model membranes, such heterogeneity is often associated with the coexistence of liquid-ordered (L_o_) and liquid-disordered (L_d_) phases, as observed in giant unilamellar vesicles (GUVs) composed of ternary mixtures of low-*T*_m_ and high-*T*_m_ lipids with cholesterol.^8^ Similar phase behavior has been reported in giant plasma membrane vesicles (GPMVs)^9^ and in yeast vacuoles *in vivo*,^10^ indicating shared biophysical principles underlying phase separation in model and biological membranes.^11^ However, microscopic phase separation in these systems is typically observed only at reduced temperatures. At physiological temperatures, membrane heterogeneity has been suggested to arise from critical fluctuations that generate transient nanoscale domains within otherwise homogeneous mixtures. ^12^

A fundamental question in membrane biophysics is which structural features of proteins dictate their preferential partitioning into one of the coexisting membrane phases. Addressing this question would, for example, shed light on the organizational principles of functional protein complexes and guide the development of novel antivirals that prevent viruses from exploiting the L_o_ domains for their entry.^13^ Since the first report of its preferential partitioning into the L_o_ phase in GPMVs,,^14^ the linker for activation of T cells (LAT) peptide, containing two cysteines available for palmitoylation, has emerged as the *bona fide* model system. Subsequent studies on GPMVs have consistently reported preferential partitioning of LAT towards the cholesterol-rich ordered regions. ^15–21^ Similar partitioning behavior has also been reported for other lipidated proteins, such as Src-family kinases,^22,23^ Ras isoforms,,^24^ and GPI-anchored proteins,,^25^ highlighting the role of post-translational modifications in modulating phase preference.

For transmembrane (TM) proteins, partitioning has been linked to transmembrane-helix properties, in particular, hydrophobic mismatch between helix length and bilayer thickness, as well as helix surface area and sequence-dependent packing.^20,21,26^ Because L_o_ membranes are thicker and more tightly packed, most TM proteins are typically found in the L_d_ phase, with only a few notable exceptions, including peripheral myelin protein 22 (PMP22),^17,18,27^ hemagglutinin,^28,29^ HIV-1 glycoprotein gp41,^30^ and perfringolysin O.^31^ Additional contributions from protein oligomerization, specific lipid–protein interactions, and membrane curvature or cytoskeletal coupling have also been reported.^32–35^

Despite these insights, a consistent picture has not yet emerged. Much of the available evidence is based on a small number of model systems, often centered around LAT, and even for individual proteins, results can be contradictory; for example, PMP22 shows different partitioning behavior in GPMVs and GUVs.^36^

On the computational side, efforts have likewise focused primarily on LAT. Coarsegrained (CG) simulations consistently report a free-energy minimum at the L_o_/L_d_ interface, reflecting a balance between the peptide’s intrinsic preference for disordered lipids and the affinity of palmitoyl anchors for ordered regions.^26,37,38^ In contrast, all-atom (AA) simulations yield less consistent results, with reported preferences ranging from L_d_ domains to interfacial localization depending on the specific system and conditions.^38,39^

While both experimental and computational studies have largely focused on a limited number of model systems, there have also been more systematic approaches to identifying general determinants of partitioning preferences using experimental datasets,^20^ bioinformatics,^40^ and machine learning.^41^ In the former approach, partitioning was found to depend on the hydrophobic mismatch between the peptide and the coexisting phases, the number of L_o_-preferring palmitoyl chains, and the interfacial tension difference of the peptide solvated in the two phases.^20^ However, this framework has significant limitations. For non-palmitoylated peptides of equal length, the interfacial tension term is the only contributor to the partitioning preference, which depends linearly on accessible surface area (ASA). As a result, chemically distinct residues contribute equally if they have similar ASA; for example, polar histidine and nonpolar leucine (both with ASA≈141 Å^2^), or charged lysine and hydrophobic methionine (both with ASA≈150 Å^2^). This coarse description neglects the underlying chemical nature of residues and their specific interactions with the lipid environment. Despite these simplifications, the model enabled predictions of raft affinity for 729 TM peptides in the human proteome, of which only a small subset showed significant preference for ordered membrane domains.^20^

Overall, our understanding of protein partitioning between L_o_ and L_d_ phases remains limited. The available data are sparse and sometimes contradictory, with reported trends varying across model systems and simulation approaches. At the same time, existing models fail to capture the influence of residue chemistry on partitioning. To address these gaps, our aim here is to clarify the physicochemical principles underlying partitioning preferences of membrane proteins by simulating a library of more than 4700 TM peptides in phase-separating lipid membranes and linking their partitioning preference to sequence- and structure-derived descriptors.

## Results

### Dataset Assembly and Characterization of Transmembrane Helix Peptides

We constructed a large, non-redundant dataset of *α*-helical transmembrane (TM) sequences from the Orientations of Proteins in Membranes (OPM) database ^42^ by combining polytopic protein and peptide entries as outlined in **Figure 1A**. Redundancy was reduced using CD-HIT^43^ clustering at 95% sequence identity, followed by transmembrane topology prediction with DeepTMHMM^44^ to extract *α*-helical TM segments. A second round of CD-HIT clustering at 90% sequence identity was then applied to the extracted helices, yielding 13,061 unique TM sequences. The length distribution was strongly dominated by 21-residue helices. To ensure comparability across sequences, we restricted the dataset to helices of this dominant length, yielding a final dataset of 4,710 sequences for all subsequent analyses. For all peptides, two lysine residues were added to each terminus to enhance stability within the bilayer and to provide a consistent interfacial environment across all sequences.

**Figure 1:**
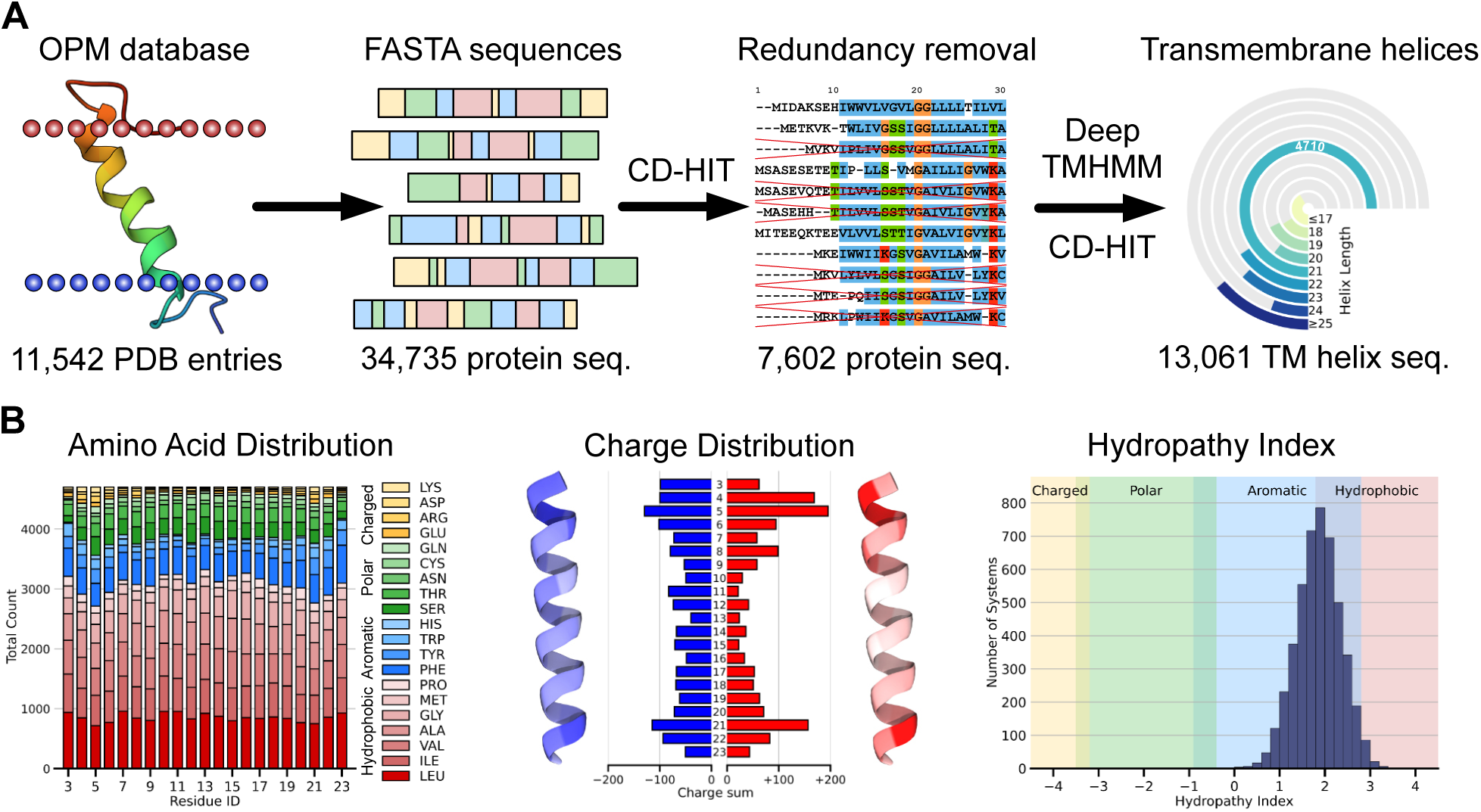
Construction and characterization of the transmembrane helix dataset. **(A)** Workflow for dataset generation. *α*-helical TM protein sequences were obtained from the OPM database and converted to FASTA format. Redundancy was reduced using CD-HIT clustering, followed by identification of TM helices with DeepTMHMM. A second clustering step yielded 13,061 unique TM helices prior to length filtering. **(B)** Sequence-based characterization of the TM helix dataset. Left: position-resolved amino acid distribution along the helix. Middle: distribution of positively (red) and negatively (blue) charged residues. Right: distribution of average hydropathy values, highlighting the predominantly hydrophobic nature of the TM helices.

As expected, the resulting TM helix dataset exhibits sequence features characteristic of membrane-spanning *α*-helices. The amino acid distribution is dominated by hydrophobic residues along the entire helix, with only minor contributions from polar and charged residues, which are mostly enriched toward the termini (**Figure 1B**, left panel). The charge distribution (middle panel) shows a clear positional asymmetry, with charged residues predominantly located near the helix ends, consistent with their preference for the membrane–water interface and the polar headgroup region, while remaining less frequent within the hydrophobic core. Finally, the hydropathy index distribution (right panel) is strongly shifted toward positive values, confirming the dataset’s overall hydrophobicity and its suitability for insertion into lipid bilayers.

### Phase Separation and Peptide Partitioning

Each of the 4,710 transmembrane helices was embedded in a rectangular lipid bilayer (12 × 8 nm^2^) composed of DPPC:DLiPC:cholesterol in a 3:3:2 ratio and simulated for 15 µs using the Martini 2.2 force field.^45–47^ This lipid composition was selected because it is known to spontaneously phase separate into coexisting liquid-ordered (L_o_) and liquid-disordered (L_d_) domains, enabling direct assessment of peptide partitioning behavior.^48,49^

Importantly, the use of the Martini 2.2 model, combined with a recently optimized cholesterol topology,^50^ ensured a physically consistent description of phase coexistence. In particular, the artificial temperature difference between coexisting phases reported in earlier simulations^51^ was largely eliminated, with the L_o_ and L_d_ phases differing by less than 1 K (**Figure 2**). This improvement avoids the formation of overly ordered and kinetically trapped domains, which have been shown to bias lipid diffusion and potentially affect protein partitioning. Consistently, lipid diffusion and mixing properties in our systems are in agreement with reference peptide-free simulations (**Figure 2**), indicating that phase behavior is not artificially enhanced.

**Figure 2:**
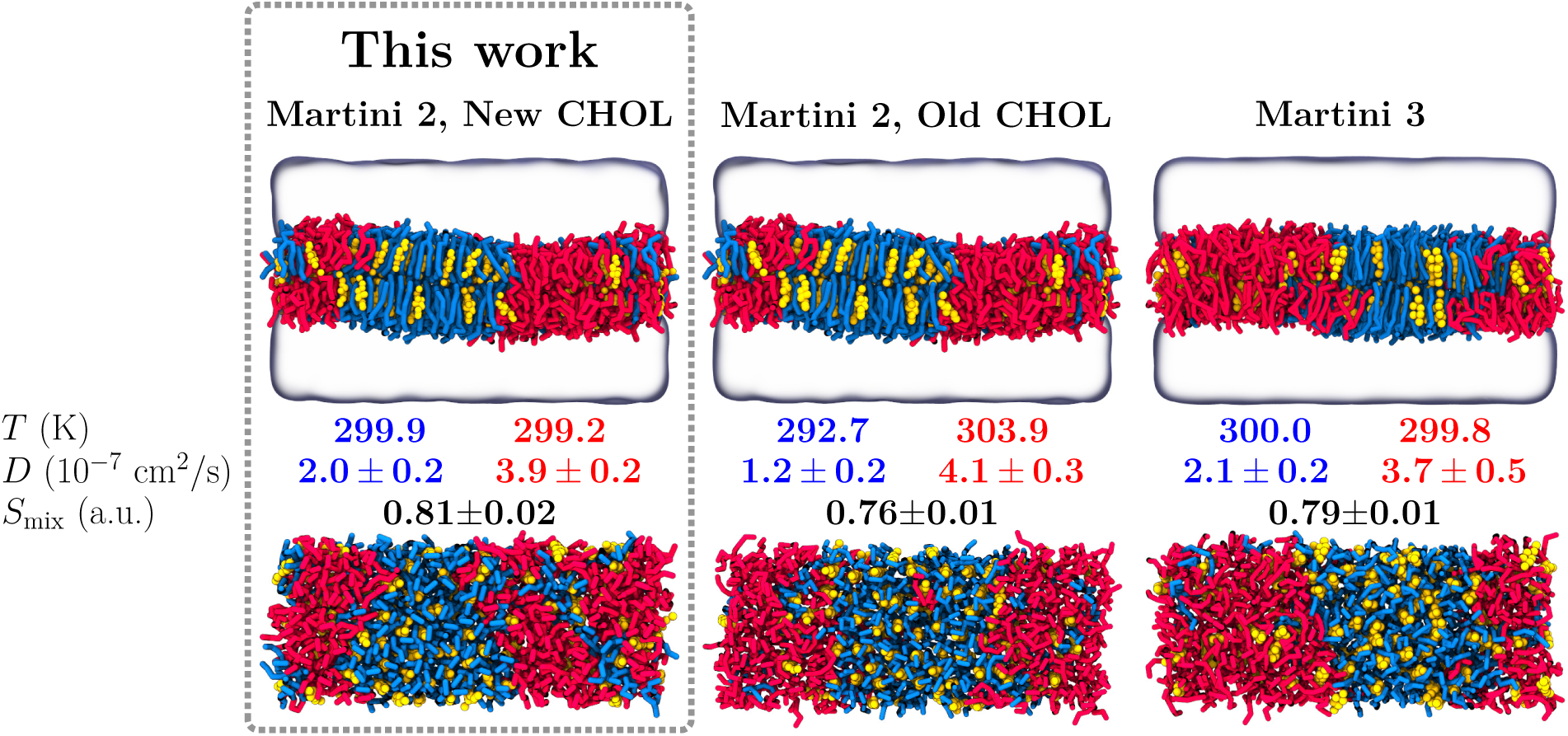
Comparison of phase behavior across Martini force fields and cholesterol models. Representative membrane configurations obtained using Martini 2.2 with the optimized cholesterol model (this work), Martini 2.2 with the original cholesterol model, and Martini 3. For each system, the temperatures of the L_d_ (blue) and L_o_ (red) phases, lipid diffusion coefficients (*D*), and mixing entropy (*S*_mix_) are reported. Snapshots illustrate the extent of domain formation in each case.

To verify the formation phase-separation domains, we quantified lipid demixing using the relative mixing entropy of neighboring lipids (see Methods for details). As shown in **Figure 3A**, the relative mixing entropy rapidly decreases within the first few microseconds and reaches a plateau after approximately 4 µs, indicating the formation of stable phase-separated domains. This is further supported by visual inspection of the membrane (inset in **Figure 3A**), where distinct cholesterol-rich L_o_ (blue) and cholesterol-poor L_d_ (red) regions are observed.

**Figure 3:**
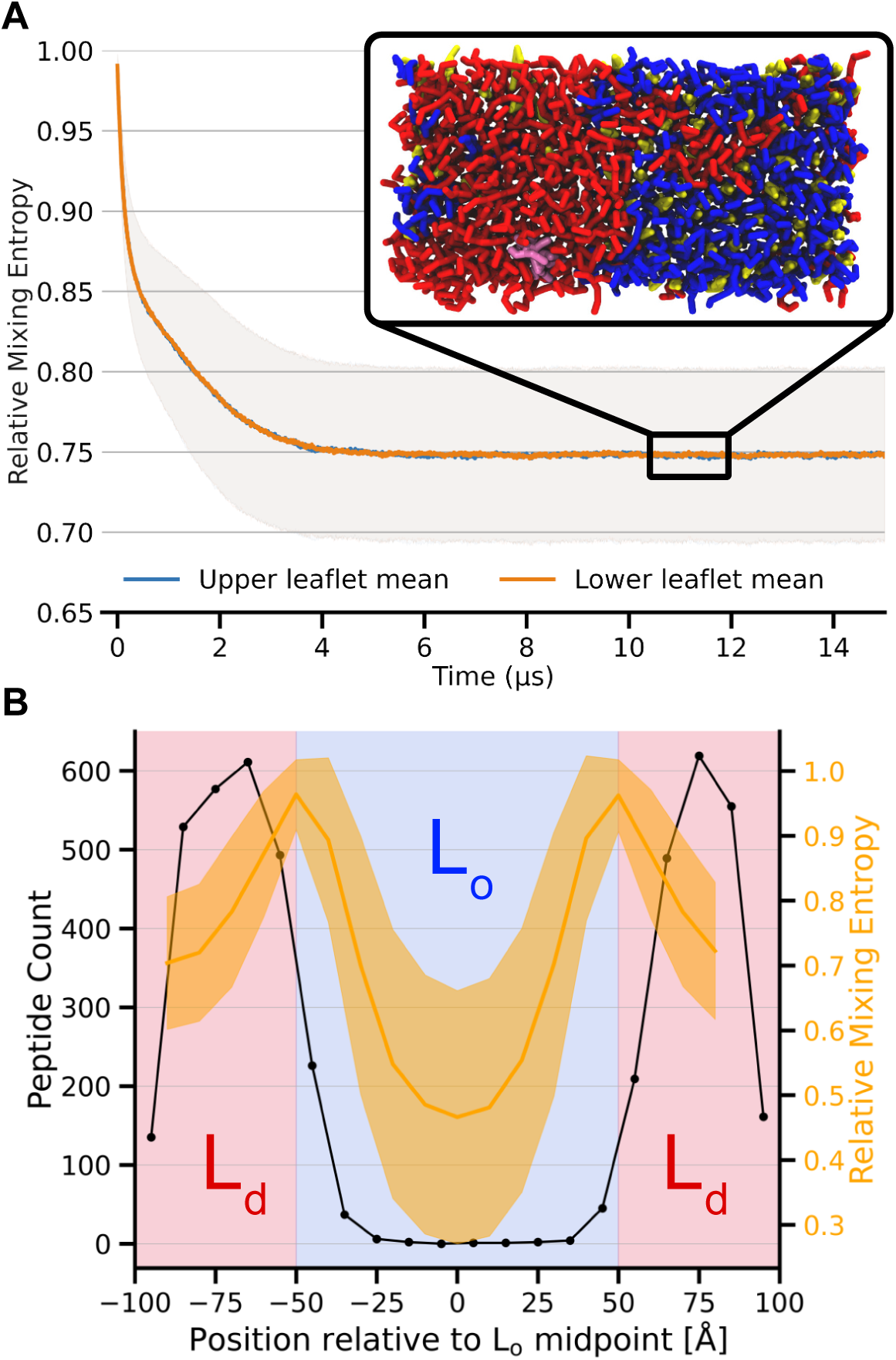
Phase separation and spatial partitioning analysis of transmembrane helices. **(A)** Time evolution of the relative mixing entropy in the upper (blue) and lower (orange) membrane leaflets. Shaded regions represent the standard deviation across 4710 simulations. The inset shows a representative final with DLiPC (red), DPPC (blue), cholesterol (yellow), and the peptide (pink), highlighting the coexistence of L_o_ and L_d_ domains. **(B)** Spatial distribution of peptides relative to the L_o_ domain midpoint. The peptide count (black) is shown together with the relative mixing entropy profile (orange, right axis). Regions corresponding to L_o_ and L_d_ phases are indicated, with the interfacial regions characterized by elevated mixing entropy.

Using the spatial profile of the relative mixing entropy across the membrane (**Figure 3B**), we identified the locations of the L_o_ and L_d_ phases as well as the interfacial regions. Analysis of peptide distributions revealed that none of the 4710 peptides preferentially partition into the L_o_ phase, while a small fraction localizes at the phase boundaries. The majority of peptides reside in the L_d_ phase, indicating a pronounced preference for the more disordered membrane environment.

### Characterization of Peptides at the Phase Interface

To further investigate the properties of peptides localized at the L_o_/L_d_ boundary, we defined an interfacial regime based on the peptide–lipid contact environment. Specifically, the fraction of contacts between each peptide and DPPC lipids was computed by considering all lipids within 12 Å of the peptide. DPPC was used as a marker of the L_o_ phase due to its fully saturated acyl chains (**Figure 4A**), which promote tight packing and preferential association with cholesterol.

**Figure 4:**
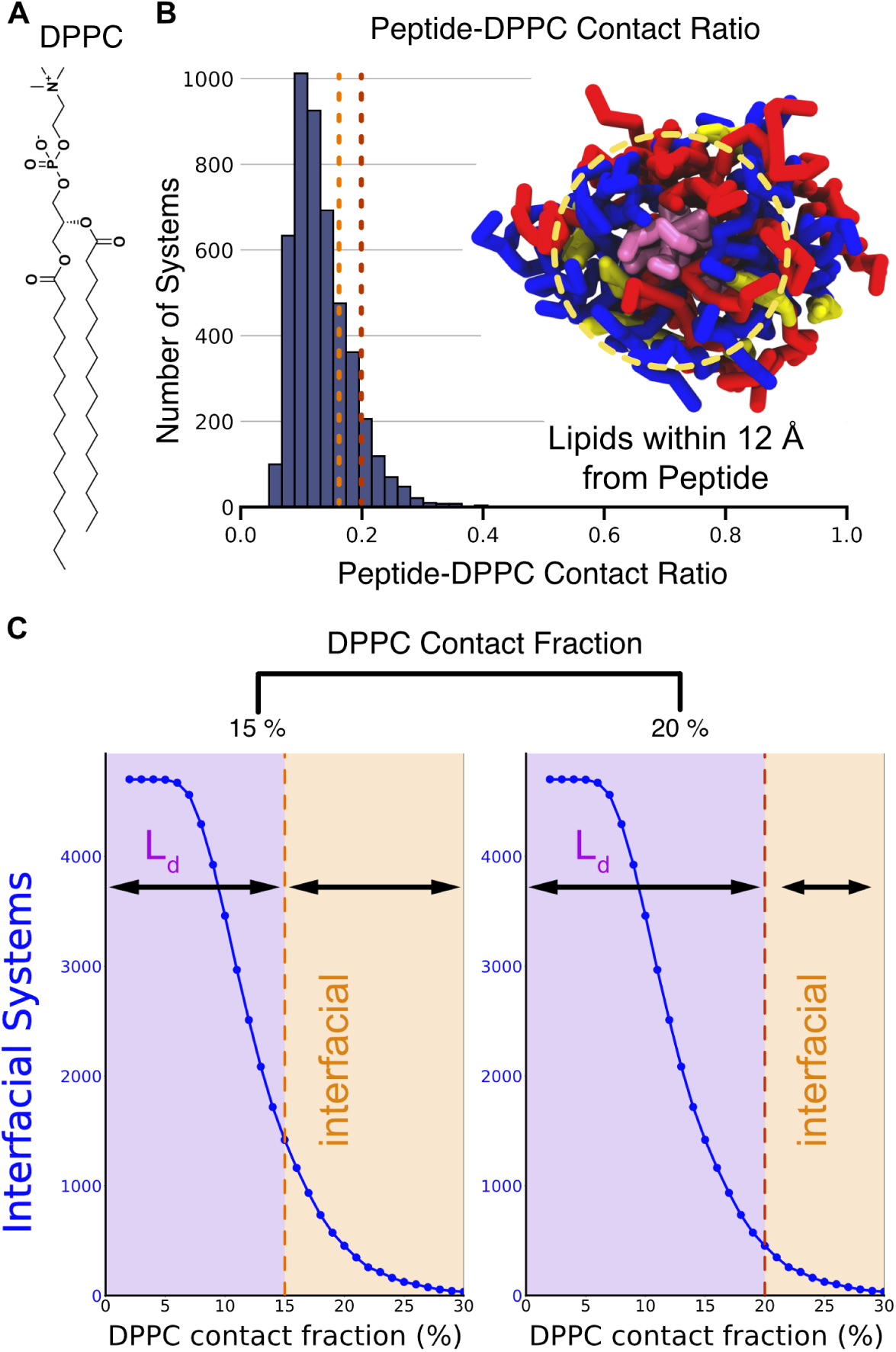
Definition of interfacial peptide partitioning. **(A)** Chemical structure of DPPC, used as an L_o_ marker to define peptide–lipid contacts. **(B)** Distribution of the peptide–DPPC contact ratio across all simulated systems, calculated from lipids within 12 Å of the peptide (illustrated on the right). Dashed vertical lines indicate threshold values used to classify peptides into L_o_ and interfacial regions. **(C)** Determination of the interfacial regime based on the DPPC contact fraction. The number of interfacial systems (blue) and statistically significant enrichments (red) are shown as a function of the DPPC contact threshold. Two representative thresholds (15% and 20%) are highlighted, illustrating the trade-off between selecting a sufficient number of interfacial peptides and maximizing statistical significance. Shaded regions indicate the L_d_ (purple) and interfacial (orange) regimes.

We employed this contact-based definition because a purely spatial definition based on the relative mixing entropy profile (**Figure 3B**) is insufficient. As also visible in the inset of **Figure 3A**, small amounts of DPPC and cholesterol can be found within the L_d_ phase, rendering a position-based identification of interfacial regions ambiguous.

As shown in **Figure 4B**, the distribution of peptide–DPPC contact ratios is strongly skewed toward low values, consistent with the overall preference of peptides for the L_d_ phase. To identify interfacial peptides, we introduced threshold values on the DPPC contact fraction. Increasing this threshold classifies fewer peptides as interfacial but reduces the statistical significance of residue enrichment (**Figure 4C**). Based on this trade-off, thresholds of 15% and 20% DPPC contact fraction were selected as representative definitions of the interfacial regime for subsequent analyses.

Using these definitions, we analyzed the amino acid composition of interfacial peptides (**Figure 5**). Compared to the remaining peptides, interfacial peptides exhibit subtle but systematic differences in residue composition across physicochemical groups. In particular, for both DPPC contact thresholds (15% and 20%), interfacial peptides show a consistent depletion of aromatic residues and a concomitant enrichment of hydrophobic residues. This trend is most pronounced for phenylalanine and tyrosine, which are less frequent at the interface, whereas aliphatic residues such as leucine, isoleucine, and valine are enriched at the interface. These trends indicate that interfacial peptides are not characterized by a single dominant residue type, but instead exhibit a combination of sequence features. Notably, the observed shifts in residue composition are consistent across both threshold definitions, supporting their robustness with respect to the precise classification of interfacial peptides. (**Figure 5A,B**).

**Figure 5:**
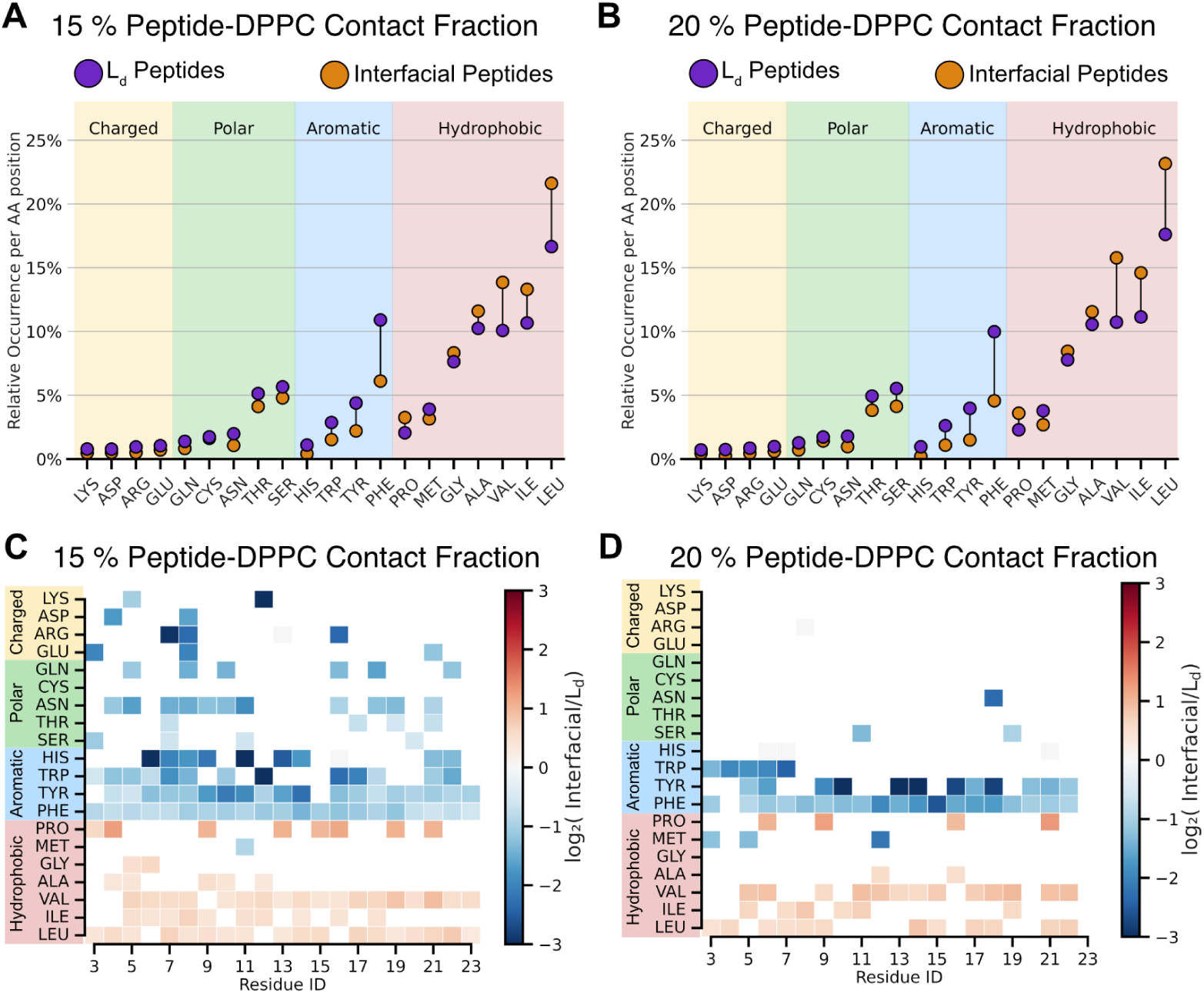
Amino acid enrichment patterns of interfacial peptides. (A,B) Relative occurrence of amino acids in interfacial peptides compared to the full dataset for two definitions of the interfacial regime based on peptide–DPPC contact fraction thresholds of 15% (A) and 20% (B). Amino acids are grouped by physicochemical properties (charged, polar, aromatic, and hydrophobic). Points represent the relative occurrence in interfacial peptides (orange) compared to the remaining peptides (purple), highlighting systematic differences in residue composition. **(C,D)** Position-resolved enrichment of amino acids in interfacial peptides, shown as log_2_(interfacial/L_d_) ratios for the 15% (C) and 20% (D) thresholds. Positive values (red) indicate enrichment in interfacial peptides, whereas negative values (blue) indicate depletion.

Position-resolved analysis further reveals that these differences are not uniformly distributed along the helix (**Figure 5C,D**). Aromatic residues (Trp, Tyr, Phe) are consistently depleted in the peptide core, with the strongest depletion observed for Trp and His at the 15% threshold and for Tyr and Phe at the 20% threshold. In contrast, hydrophobic residues such as Leu, Ile, and Val show enrichment across much of the helix, indicating a general preference for aliphatic side chains in interfacial peptides. Polar and charged residues are likewise depleted in the core region, although this trend is more pronounced at the 15% threshold and less consistent at 20%, indicating some sensitivity to the definition of the interfacial regime. Among all residues, proline exhibits the strongest enrichment; however, this effect is localized to specific positions rather than distributed uniformly along the helix. Overall, these results show that peptides residing at the L_o_/L_d_ interface possess distinct sequence characteristics compared to peptides embedded in the L_d_ phase. These differences are reflected in position-dependent shifts in residue composition, indicating that both amino acid identity and spatial distribution contribute to interfacial localization. The observed trends are largely consistent across threshold definitions, although the magnitude of enrichment and depletion varies. However, the apparent positional specificity may be influenced by correlations between residues within individual sequences, which are not explicitly accounted for in this analysis.

## Discussion

In this work, we adopted a high-throughput computational approach to investigate the determinants of transmembrane helix partitioning in phase-separating membranes. In contrast to previous studies, which have largely focused on one or a few model peptides, our analysis of more than 4,700 sequences enables the identification of statistically significant trends and provides a broader, data-driven perspective on membrane partitioning.

A key technical aspect of this study is the use of an improved cholesterol model, which eliminates the artificial temperature difference between coexisting phases reported in earlier Martini simulations.^51^ This artifact has been shown to exaggerate phase separation and reduce lipid mobility, rendering the L_o_ phase almost gel-like and thus potentially biasing the partitioning behavior of embedded proteins.

By mitigating this issue, our simulations provide a more balanced representation of L_o_/L_d_ coexistence. In this context, it is notable that previous coarse-grained studies frequently reported strong interfacial localization or even stabilization of ordered domains by peptides,^26,37^ whereas our results suggest a weaker coupling between proteins and the ordered phase. This indicates that some conclusions drawn from earlier coarse-grained simulations may reflect model-specific artifacts rather than intrinsic physicochemical preferences.

Across all analyzed systems, we did not observe a single peptide with a clear preference for the L_o_ phase. This finding is consistent with the strong preference of transmembrane helices for the L_d_ phase reported in both experiments and simulations of simplified model membranes.^20,36^ The thicker and more tightly packed nature of the L_o_ phase imposes an energetic penalty for accommodating transmembrane helices, an effect commonly attributed to hydrophobic mismatch and steric constraints. Nevertheless, because 21-residue helices are among the most prevalent transmembrane helix lengths in membrane proteins, the peptides considered here remain biologically relevant in length. Curiously, our capped peptides are ≈3.8 nm long, and equally long KKL_21_KK-peptides were recently found to spontaneously partition to membrane thicknesses of 3.8±0.6 nm and demonstrate a tilt of 26±12*^◦^*.^52^ This indicates that hydrophobic mismatch alone is not sufficient to explain the observed tendency. We simulated single-phase L_d_ (100% DIPC) and L_o_ (60% DPPC & 40% CHOL)) systems, and their thicknesses of 3.6 nm and 4.4 nm are within this spontaneously accessible range, with zero mismatch actually occurring for peptides at the phase boundary. As L_d_-partitioning is dominant for our large and diverse sequence space, we can conclude that the steric effects play a major role over sequence-specific variations.

At the same time, our findings highlight an important discrepancy with observations in more complex systems such as GPMVs and living cells, where several membrane proteins have been reported to partition into ordered domains.^14,15^ This discrepancy highlights a limitation of simplified ternary mixtures, which do not fully capture the physicochemical environment relevant for lipid raft-associated protein sorting. In biological membranes, additional factors such as lipidation, protein oligomerization, specific lipid–protein interactions, and the crowded cellular environment likely contribute to stabilizing localization in ordered regions.^4,5^ Moreover, the phases are more similar in GMPVs than in synthetic vesicles, as revealed by generalized polarization and cryo-ET.^11,53,54^ Our results, therefore, provide a baseline for the intrinsic partitioning behavior of isolated transmembrane helices, against which these additional contributions can be evaluated.

Despite the absence of L_o_-preferring peptides, our analysis revealed a subset of sequences that preferentially localize at the L_o_/L_d_ interface. These interfacial peptides exhibit distinct sequence characteristics and may act as line-active agents (“linactants”) that lower the line tension, analogous to a surfactant lowering surface tension, thereby leading to boundary destabilization and changes in domain morphology. Such behavior, reported for certain lipids,^55,56^ could have important implications for membrane organization, for example by reducing line tension or stabilizing nanoscale domains. This behavior is consistent with the notion that the phase boundary represents a distinct environment that can be selectively stabilized by specific sequence features.

Finally, our results place recent attempts to predict raft affinity from sequence features into context.^20,41^ While such approaches capture general trends, our analysis highlights the limitations of coarse descriptors such as accessible surface area. In particular, both residue identity and positional organization contribute to partitioning behavior, indicating that more detailed representations of sequence chemistry are required for accurate predictions.

Overall, our findings indicate that intrinsic partitioning of transmembrane helices is strongly biased toward the disordered phase, with interfacial localization emerging as a secondary but well-defined behavior. This provides a baseline for sequence-dependent partitioning and underscores that localization to ordered domains in biological membranes likely requires additional contributions beyond the transmembrane segment. The functional consequences of this interfacial localization, and its potential role in regulating membrane heterogeneity, remain to be explored and will be the focus of future work.

## Methods

### Construction of the Transmembrane Helix Sequence Dataset

*α*–helical transmembrane protein sequences were collected from the Orientations of Proteins in Membranes (OPM) database (October 2024 release).^42^ Two OPM subsets were combined: OPM alpha-helical polytopic (8,078 entries) and OPM peptides (8,915 entries), yielding a total of 16,993 PDB ID entries, including duplicate identifiers. These entries were merged into a single file (OPM combined.txt). Duplicate PDB identifiers were removed, resulting in 11,542 unique PDB IDs. For each unique PDB ID, protein sequences were retrieved from the PDB in FASTA format using an automated script (pdbID2fasta.sh), producing 34,735 protein sequences (OPM fasta.fasta). To reduce redundancy prior to transmembrane helix prediction, sequences were clustered using Cluster Database at High Identity with Tolerance (CD-HIT)^43^ at 95% sequence identity (-c 0.95, -n 5). This step yielded 7,602 non-redundant protein sequences (OPM clustered 0.95.fasta). Transmembrane helices were then identified using DeepTMHMM^44^ version 1.0.42 (via BioLib: https://github.com/donovan-h-parks/biolib), which predicts transmembrane topology based on deep learning. All predicted *α*-helical transmembrane segments were extracted, resulting in 17,220 transmembrane helix sequences (TMhelixes.fasta). To further remove sequence redundancy among transmembrane helices, a second clustering step was performed using CD-HIT at 90% sequence identity (-c 0.9, -n 5). This final filtering step yielded 13,061 unique transmembrane helix sequences (TMhelixes clustered 0.90.fasta). Among these, 21–amino acid–long helices dominated the length distribution, with 4,710 unique sequences (TMhelixes clustered 0.90 21AA.fasta), which constitute the dataset used for all subsequent analyses. All datasets and scripts generated in this study are publicly available on Zenodo under DOI: https://doi.org/10.5281/zenodo.17967865.

### Molecular Dynamics Simulations

#### System setup

The 4,710 sequences of 21 amino acids in length were converted into idealized three-dimensional *α*-helical peptides using an in-house Python script (fasta2helix v0.3.py), based on the PeptideBuilder library.^57^ These helical peptides were capped on both ends by two lysines, coarsegrained^58^ and inserted in a membrane patch^59^ with lateral dimensions of 12×8 nm^2^ and a composition of 3:3:2 of dilinoleoylphosphatidylcholine (DLiPC, often referred to as DUPC or DIPC in simulation studies):dipalmitoylphosphatidylcholine (DPPC):cholesterol (CHOL). Pure DLiPC and DPPC:Cholesterol mixtures have thicknesses of 3.6 nm and 4.4 nm in our simulations. Equally long peptides were found to favourably partition to membranes with a thickness of 3.8 nm in our recent unbiased simulations, whereas values from 3.7 nm to 4.4 nm were achievable within thermal fluctuations.^52^ This indicates that mismatch is not a major contributor in partitioning free energy, especially compared to the reported total partitioning free energies.^20,26,38,39^ The dimension normal to the membrane was set to 10 nm, with water and 10% antifreeze particles used as the solvent. Any positive charge in the peptide (mostly arising from the Lys capping residues) was accounted for by Cl*^−^* ions. The systems were simulated for 15 µs each (≈71 ms of total simulation time), during which the initially random lipid mixture spontaneously separated into coexisting L_o_ and L_d_ phases, and the peptide partitioned into its preferred phase or at their boundary.

#### Simulations parameters

We modeled the peptide and lipids using the Martini 2.2 model,, ^45–47^ which captures the spontaneous demixing of lipids into the L_o_ and L_d_ phases.^49^ This choice was motivated by the TM orientation being unstable for many peptides^60^ in the more recent Martini 3 model,^61,62^ and since we observed no differences in the phase coexistence between the two versions. In the peptide-free system, the degree of phase separation, quantified by the mixing entropy, was similar for the two models. Additionally, the diffusion coefficients extracted for the low-*T*_m_ and high-*T*_m_ lipids agreed between them (Figure 2).

We also used the recently optimized cholesterol topology,,^50^ which eliminated the temperature difference of the two phases ^51^ present with the old cholesterol. In our simulations, we observed the temperatures of the low-*T*_m_ and high-*T*_m_ lipids to differ by a mere 0.7 K, compared to 11.2 K observed with the old cholesterol model. This artifact manifested itself in overly-demixed membrane and a low diffusion of the low-*T*_m_ lipids (Figure 2), and likely played a major role in earlier studies’ finding that no peptides prefer the overly ordered and stiff L_o_ phase.

The exaggerated oligomerization of membrane proteins in Martini 2.2^63^ was not an issue as all studied systems only contained a single peptide. All simulations were performed in the NPT ensemble using GROMACS with a time step of 25 fs. Production runs were propagated for 400 M integration steps, corresponding to 15 µs per system. Compressed coordinates and energies were saved every 10 ns. Neighbor lists were updated every 20 steps using the Verlet scheme under periodic boundary conditions in all directions. A neighbor-list cutoff of 1.35 nm was used together with a negative Verlet buffer tolerance (−1), ensuring stable membrane fluctuations and avoiding neighbor-list artifacts.^64^

Electrostatic interactions were treated using the reaction-field method ^65^ with a cutoff of 1.1 nm, a relative dielectric constant of 15, and *ɛ*_rf_ = 0. Van der Waals interactions were truncated at 1.1 nm using the potential-shift-Verlet scheme. Temperature was maintained at 300 K using the velocity-rescale thermostat^66^ (*τ_t_* = 1.0 ps), with separate coupling groups for membrane and solvent. Pressure was controlled semi-isotropically at 1 bar using the c-rescale barostat^67^) (*τ_p_* = 4.0 ps; compressibility = 3 × 10*^−^*^4^ bar*^−^*^1^).

All simulations were performed on AMD Instinct MI250X accelerators^68^ on the LUMI supercomputer. The simulation outputs are available on Zenodo at DOI: https://doi.org/10.5281/zenodo.17967865.

### Analysis

#### System Preparation and Filtering

Of the 4,710 peptides, 7 were removed from further analysis as their center of mass (COM) left the hydrophobic membrane core, leaving 4,703 peptides for analysis. Residues 1, 2, 24, and 25 were excluded, as they correspond to lysine residues added to anchor the peptides in the membrane. To reduce computational cost, water and ions surrounding the membrane were removed prior to analysis.

#### Sequence-Based Descriptors

General properties of the peptides, including amino acid composition, charge distribution, and average hydropathy (based on the Kyte–Doolittle scale^69^), were calculated directly from the sequence.

#### Peptide Orientation Analysis

To assess the stability of transmembrane insertion, the tilt angle of each peptide was calculated for every frame of the last 5 *µ*s of the trajectory. The angle was defined as the arccosine of the angle between the membrane normal (z-axis) and the vector connecting the backbone atoms of the N- and C-terminal lysines. Simulations with tilt angles greater than 65*^◦^* were visually inspected to determine whether the peptide had left the membrane. This led to the exclusion of 6 of the 7 systems that had not been captured earlier.

#### Quantification of Phase Separation

Phase separation was quantified using the mixing entropy based on conditional probabilities as introduced by Brandani et al.:^70^

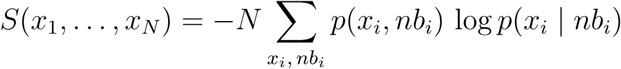

where *p*(*x_i_, nb_i_*) is the joint probability of finding lipid types *x_i_* and *nb_i_* as neighbors, and *p*(*x_i_* | *nb_i_*) is the corresponding conditional probability. Lipid neighborhoods were defined based on distances between headgroup beads (PO4 for DLiPC and DPPC, ROH for cholesterol), with the nearest neighbor identified for each lipid. Probabilities were normalized such that the total probability equals 1.

The mixing entropy was normalized using the theoretical maximum:

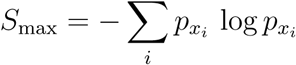

yielding a normalized entropy *S* = 1.

#### Spatial Alignment and Domain Identification

Bilayers were aligned such that L_o_ domains overlapped in the *x*–*y* plane. The final simulation frame was used for analysis, assuming that domain interfaces primarily translate along the *y* axis. The simulation box was divided into 18 stripes of 1 nm width along the *x* axis, and the mixing entropy was computed for each stripe and normalized by its local maximum. Stripes with the highest entropy were identified as interfacial regions. The two interfaces were determined by pairing the stripe with the highest entropy and its neighbors with the stripe located furthest away and its neighbors. The domain with higher DPPC density was assigned as L_o_.

### Peptide Partitioning Analysis

Two complementary approaches were used to classify peptide partitioning.

#### Position-based classification

Peptide positions were expressed relative to the L_o_ midpoint. Boundaries between L_o_, interfacial, and L_d_ regions were defined based on the mixing entropy profile.

#### Contact-based classification

For each frame of the last 5 *µ*s, lipids within 1.2 nm of the peptide were identified. The DPPC contact ratio was defined as:

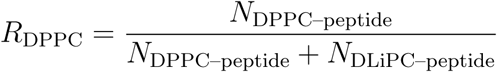

Peptides were classified based on *R*_DPPC_ using two threshold definitions:

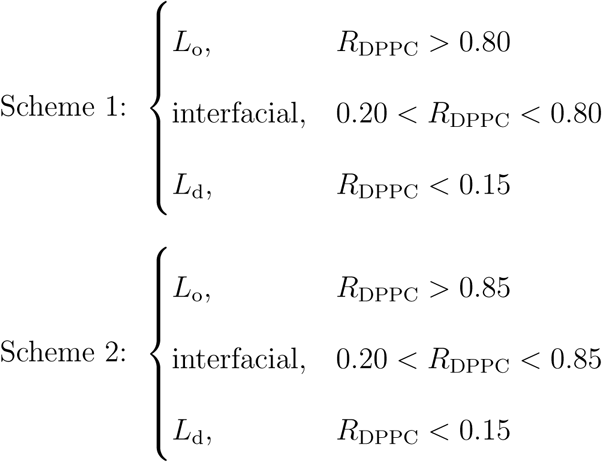

Both schemes were further analyzed and compared.

### Statistical Analysis

Amino acid distributions were compared between environments by computing probability ratios. Statistically significant enrichments were identified using Fisher’s exact test.

## Acknowledgement

This work is supported by grants of the Deutsche Forschungsgemeinschaft to FL (SFB-1638/1 – 511488495 - Z01). MJ thanks The Finnish Academy of Science and Letters for the Väisälä project grant and CSC–IT Center for Science for computational resources (LUMI Extreme Scale project “PepMemML: Design of optimal peptide sequences for membrane-related applications using high-throughput molecular simulations and machine learning”). FL acknowledges the computing resources provided by the state of Baden-Württemberg through bwHPC and the German Research Foundation (DFG) through grant INST 35/1597-1 FUGG. FL gratefully acknowledges the data storage service SDS@hd supported by the Ministry of Science, Research and the Arts Baden-Württemberg (MWK) and the German Research Foundation (DFG) through grant INST 35/1503-1 FUGG.

